# Causal role of L-glutamine in sickle cell disease painful crises: a Mendelian randomization analysis

**DOI:** 10.1101/872358

**Authors:** Yann Iboudo, Melanie E. Garrett, Pablo Bartolucci, Carlo Brugnara, Clary B. Clish, Joel N. Hirschhorn, Frédéric Galactéros, Allison E. Ashley-Koch, Marilyn J. Telen, Guillaume Lettre

## Abstract

In a recent clinical trial, the metabolite L-glutamine was shown to reduce painful crises in sickle cell disease (SCD) patients. To confirm this observation and identify other metabolites implicated in SCD clinical heterogeneity, we profiled 129 metabolites in the plasma of 705 SCD patients. We tested correlations between metabolite levels and six SCD-related complications (painful crises, cholecystectomy, retinopathy, leg ulcer, priapism, aseptic necrosis) or estimated glomerular filtration rate (eGFR), and used Mendelian randomization (MR) to assess causality. We found a causal relationship between L-glutamine levels and painful crises (N=1,278, odds ratio (OR) [95% confidence interval] = 0.68 [0.52 – 0.89], *P*=0.0048). In two smaller SCD cohorts (N=299 and 406), the protective effect of L-glutamine was observed (OR=0.82 [0.50-1.34]), although the MR result was not significant (*P*=0.44). We identified 66 significant correlations between the levels of other metabolites and SCD-related complications or eGFR. We tested these correlations for causality using MR analyses and found no significant causal relationship. The baseline levels of quinolinic acid was associated with prospectively ascertained survival in SCD patients, and this effect was dependent on eGFR. Metabolomics provide a promising approach to prioritize small molecules that may serve as biomarkers or drug targets in SCD.

## INTRODUCTION

Sickle cell disease (SCD) is one of the most common Mendelian diseases in the world, affecting millions of patients living in Sub-Saharan Africa and the Indian sub-continent (1). In the United States, >100,000 individuals, mostly of African descent, live with SCD, and healthcare costs associated with SCD management and treatment are substantial (2). Although fundamentally a disease of the blood – caused by mutations in the β-globin gene *HBB* – SCD is characterized by systemic and debilitating complications, such as painful crises, stroke, pulmonary hypertension and kidney failure. Unfortunately, there are no robust prognostic biomarkers to predict who will develop which complications, and when. SCD treatment still relies primarily on chronic blood transfusions and hydroxyurea (HU), a drug that acts partly by raising the concentration of antisickling fetal hemoglobin (HbF)(3).

Progress in gene therapies and genome editing technologies now offer realistic hope of developing a cure for SCD (4). However, these complex clinical interventions are unlikely to benefit most SCD patients worldwide in the short term. Therefore, we need to continue searching for novel biomarkers and drug targets for SCD. Recently, the US Food and Drug Administration approved a second molecule, L-glutamine, to treat SCD. In a double-blind phase 3 clinical trial, L-glutamine was shown to reduce the number of painful crises over a 48-week period (5). The emergence of L-glutamine as a therapy was based on decades of work investigating the role of oxidative stress in SCD pathophysiology (6). Red blood cells (RBC) from SCD patients have high oxidative stress and a compromised ability to counteract free radicals due to a low ratio of the reduction-oxidation (redox) co-factor nicotinamide adenine dinucleotide (NAD) and its reduced form ([NADH]:[NAD^+^+NADH])(7). L-glutamine is one of the most abundant amino acids in the human body and in addition to its role in protein synthesis, is required to synthesize NAD. Treatment with L-glutamine increases the NAD redox ratio and reduces adhesion of sickle RBC to endothelial cells, a hallmark of vaso-occlusive painful crises (8, 9).

Metabolites, like L-glutamine, are small molecules (e.g. amino acids, sugars, lipids) that result from the activities of endogenous enzymes (10). The development of high-throughput mass spectrometry-based methodologies makes it possible to profile 100-1000s of metabolites in human biospecimens. Such metabolomic studies have been used to identify metabolite signatures of diseases, but also to pinpoint specific metabolites that may have prognostic and/or therapeutic values (11, 12). Metabolite levels are variable between individuals (in disease, but also in health) and large genetic studies – termed metabolite genome-wide association studies (mGWAS) – have identified 1000s of genetic variants that control them. Besides providing an opportunity to characterize the biological pathways that control metabolite levels, these genetic discoveries become powerful instruments for Mendelian randomization (MR) studies. MR uses genetic variants to determine the effect of genetically modulated phenotypes on disease outcome (13). MR mimics randomized clinical trial as it harnesses the random allocation of parental alleles when they are passed on to their offspring. As a consequence, the alleles are independently distributed in the population and free from potential confounders (14, 15). Previous MR studies have validated many drug targets for various human diseases (e.g. statins that lower LDL-cholesterol levels to reduce coronary artery disease (CAD) risk), but have also been useful to rule out many biomarkers as potential causal factors (e.g. HDL-cholesterol or C-reactive protein for CAD)(16–18).

In SCD, only a limited number of studies have used metabolomic approaches to tackle clinical heterogeneity. Zhang *et al*. discovered increased adenosine levels in blood from SCD patients and transgenic mice: they showed that higher adenosine levels exacerbated sickling, hemolysis and organ damage (19). Additionally, the same group found that sphingosine-1-phosphate (S1P) and 2,3-bisphosphoglycerate (2,3-BPG) blood concentrations are elevated in SCD patients and mice, which results in the re-programming of the glycolysis program and enhanced disease severity (20, 21). Finally, Dargouth *et al*. profiled the metabolome of RBC from healthy individuals and SCD patients and identified several metabolites that highlight differences between the two groups in glycolysis, membrane turnover, and glutathione and nitric oxide metabolism(22). Although exciting, these pioneering metabolomic studies were performed in a limited number of SCD patients (N=14-30) and did not take advantage of MR methodology to address causality.

To prioritize metabolites that may be important biomarkers or drug targets for SCD, we profiled 129 known metabolites, including L-glutamine, in the plasma of 705 SCD patients. First, we used MR to test the causal relationship between L-glutamine and painful crises. Second, we tested the association between all measured metabolites and SCD-related complications and combined these results with previous mGWAS findings to perform MR studies. Finally, we tested if baseline plasma metabolite levels were associated with survival in our SCD cohorts. Our results highlight the value of combining genetic and metabolomic strategies to disentangle the complex pathophysiology of SCD.

## SUBJECTS AND METHODS

### Study participants

Sample collections and procedures were in accordance with the institutional and national ethical standards of the responsible committees and proper informed written consent was obtained. The Genetic Modifier (GEN-MOD), the Cooperative Study of Sickle Cell Disease (CSSCD), and the Duke University Outcome Modifying Genes (OMG) cohorts have been described elsewhere (23–25). In particular for GEN-MOD, a dedicated research assistant validated all clinical information. Demographic and clinical information for each SCD cohort is available in **Table 1**.

**Table 1.** Demographics and clinical information. Sickle cell disease patients from three cohorts were included in this study. For the CSSCD, all data are prospective and we only considered patients with genome-wide genotyping data available. For GEN-MOD and OMG, all data were collected at baseline and are retrospective, except survival which is prospective. ^1^Painful crises in GEN-MOD and OMG are defined as crises requiring hospitalization which was dichotomized (individuals with ≥1 painful crises in the last 12 months are assigned as cases, while individuals with no painful crisis are assigned as controls). In the CSSCD, painful crises are defined as painful episodes requiring emergency room visits, and we dichotomized the data as no crisis (control) or at least one crisis (case) during the follow-up period. For all quantitative variable, we provide mean ± standard deviation. LDH, lactate dehydrogenase; RBC, red blood cell; MCH: mean corpuscular hemoglobin; MCV: mean corpuscular volume; eGFR, estimated glomerular filtration rate; NA, not available.

| Characteristic | GEN-MOD | OMG | CSSCD |
| --- | --- | --- | --- |
| Sex (male/female) | 222/184 | 163/136 | 616/662 |
| Age (year) | 31 ± 9 | 35 ± 14 | 14 ± 12 |
| β-globin genotypes (HbSS/<br>HbS $\beta^0$-thal/ HbSS α-thal/ HbSC/ HbS $\beta^+$) | 406/0/0/0/0 | 255/12/0/23/9 | 883/0/395/0/0 |
| Painful crises (cases/controls)<sup>1</sup> | 150/180 | 161/128 | 194/907 |
| Leg ulcer (cases/controls) | 30/300 | 79/214 | 185/970 |
| Cholecystectomy (cases/controls) | 200/206 | 173/72 | 152/932 |
| Aseptic necrosis (cases/controls) | 94/236 | 93/194 | 164/991 |
| Priapism (cases/controls) | 41/116 | 55/236 | 96/460 |
| SCD retinopathy (cases/controls) | 182/67 | 65/210 | 274/292 |
| SCD survival (cases/controls) | 19/384 | 35/91 | 44/1235 |
| Bilirubin (mg/dL) | 3.5 ± 2.1 | 2.9 ± 2.4 | 3.3 ± 2.2 |
| eGFR (mL/min<br>per 1.172 m<sup>2</sup>) | 143.6 ± 22.8 | 126.0 ± 40.3 | 165.3 ± 47.0 |
| Hemoglobin (g/dL) | 8.8 ± 1.3 | 8.2 ± 1.8 | 8.4 ± 1.3 |
| Hematocrit (%) | 25.8 ± 4.5 | 25.3 ± 5.7 | 24.8 ± 4.03 |
| Lactate dehydrogenase (units/L) | 400.4 ± 144.7 | 326.8 ± 240.2 | 451.6 ± 244.7 |
| MCH (pg) | $29.3 \pm 4.1$ | $32.0 \pm 4.8$ | $29.8 \pm 3.1$ |
| MCV (fL) | $87.0 \pm 10.2$ | $92.2 \pm 12.7$ | $89.2 \pm 8.5$ |
| RBC count ($\times 10^6$ cells/μL) | $3.0 \pm 0.80$ | $2.9 \pm 0.80$ | $2.8 \pm 0.57$ |

### Metabolomics Profiling

Plasma metabolites were profiled using two complimentary liquid chromatography tandem mass spectrometry (LC-MS) methods. Amino acids, amino acid metabolites, acylcarnitines, and other cationic polar metabolites were measured using a Nexera X2 U-HPLC (Shimadzu Corp.) coupled to a Q Exactive Hybrid Quadrupole Orbitrap Mass Spectrometer (Thermo Fisher Scientific). Plasma samples (10 μl) were prepared via protein precipitation, with the addition of 9 volumes of acetonitrile/methanol/formic acid (74.9:24.9:0.2; v/v/v) containing stable isotope-labeled, quality control internal standards (valine-d8, Sigma-Aldrich; St. Louis, MO; and phenylalanine-d8, Cambridge Isotope Laboratories; Andover, MA). The samples were centrifuged (10 min, 9,000 x g, 4°C), and the supernatants were injected directly onto a 150 x 2 mm, 3 μm Atlantis HILIC column (Waters). The column was eluted isocratically at a flow rate of 250 μL/min with 5% mobile phase A (10 mM ammonium formate and 0.1% formic acid in water) for 0.5 minute followed by a linear gradient to 40% mobile phase B (acetonitrile with 0.1% formic acid) over 10 minutes. MS analyses were carried out using electrospray ionization in the positive ion mode using full scan analysis over 70-800 *m/z* at 70,000 resolution and 3 Hz data acquisition rate. Other MS settings were: sheath gas 40, sweep gas 2, spray voltage 3.5 kV, capillary temperature 350°C, S-lens RF 40, heater temperature 300°C, microscans 1, automatic gain control target 1e6, and maximum ion time 250 ms. Raw data were processed using TraceFinder software (Thermo Fisher Scientific; Waltham, MA) for supervised, targeted extraction of data from a subset of lipids and Progenesis QI (Nonlinear Dynamics; Newcastle upon Tyne, UK). Organic acids, sugars, purines, pyrimidines, and other anionic polar metabolites were measured using an ACQUITY UPLC (Waters Corp, Milford MA) coupled to a 5500 QTRAP triple quadrupole mass spectrometer (AB SCIEX, Framingham MA). Plasma samples (30 μL) were extracted using 120 μL of 80% methanol containing 0.05 ng/μL inosine-15N4, 0.05 ng/μL thymine-d4, and 0.1 ng/μL glycocholate-d4 as quality control internal standards (Cambridge Isotope Laboratories, Inc., Tewksbury MA). The samples were centrifuged (10 min, 9,000 x g, 4°C) and the supernatants (10 μL) were injected directly onto a 150 x 2.0 mm Luna NH2 column (Phenomenex, Torrance CA). The column was eluted at a flow rate of 400 μL/min with initial conditions of 10% mobile phase A (20 mM ammonium acetate and 20 mM ammonium hydroxide (Sigma-Aldrich) in water (VWR)) and 90% mobile phase B (10 mM ammonium hydroxide in 75:25 v/v acetonitrile/methanol (VWR)) followed by a 10 min linear gradient to 100% mobile phase A. The ion spray voltage was −4.5 kV, the source temperature was 500°C, and multiple reaction monitoring (MRM) settings for each metabolite were determined using authentic reference standards. Raw data were processed and visually reviewed using MultiQuant software (AB SCIEX, Framingham MA).

### Metabolomics pre-processing

We removed metabolites with >20% missing values. We imputed missing metabolite values using the k-nearest neighbors algorithm (26) as implemented in the R package *impute*. We log_10_-transformed metabolite values, and applied batch effect correction based on metabolites’ dates of extraction, the types of ionization (negative and positive ionization), and whether they were obtained from targeted or untargeted approaches. Finally, we applied batch effect correction based on the year of profiling, since sample collection occurred within a 3 years span (2015-2017). We conducted all batch effect correction using combat (27). Using a linear model, we then derived residuals correcting for age, sex, SCD genotypes, and HU usage. **Figure 1** summarizes the design of our metabolomic experiment. Although we captured many unknown metabolites, which we used as part of the quality-control steps, this study focuses on the 129 known metabolites that were available in both GEN-MOD and OMG.

**Figure 1.**
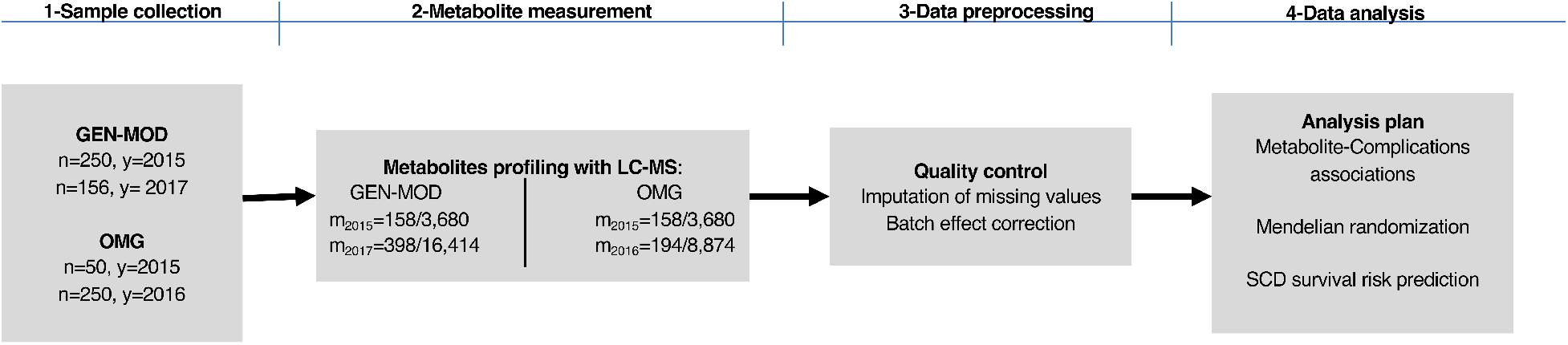
Study design of the metabolomic study in sickle cell disease (SCD) patients. 250 GEN-MOD samples and 50 OMG samples were profiled in 2015, 250 OMG samples were profiled in 2016, and 156 GEN-MOD samples were profiled in 2017. Known/targeted and unknow/untargeted metabolites were measured using liquid-chromatography in tandem with mass spectrometry (LC-MS). Data preprocessing involved standard quality-control procedures, imputation of missing values, batch-effect correction and data scaling. Data analysis included association testing of known metabolites with SCD-related complications, Mendelian randomization, and SCD survival prediction using statistical modelling. n, number of patients included in the study; y, year during which metabolites were measured; m, number of targeted/untargeted metabolites measured in each year.

### Pairwise association between metabolite levels and SCD complications or survival in GEN-MOD and OMG

To test the association between metabolite levels and SCD complications (painful crises, aseptic necrosis, cholecystectomy (gall bladder removal), retinopathy, priapism, leg ulcer, survival), estimated glomerular filtration rate (eGFR, calculated using the chronic kidney disease epidemiology collaboration (CKD-EPI) equation (28)) or to predict the risk of prospectively ascertained death (survival), we implemented a permutation procedure that considers the correlation between metabolite levels. We randomly permuted the phenotype of interest and computed 100,000 P-values (for each metabolite) in a linear or a logistic model. We then stored the smallest P-value out of the 100,000, and obtained the adjusted/permutated P-value (*P*_perm_) by comparing the number of times the permutated P-values are smaller than the observed P-values:

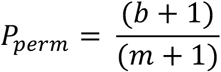

where *b* is the number of times *P*_perm_ is greater or equal than *P*_obs_, and *m* the number of permutations. The procedure was implemented in the R statistical package.

### Genetic association study in the CSSCD

DNA genotyping and genotype imputation in the CSSCD have been described in detail elsewhere (25). We restricted our analysis to markers with imputation quality r^2^ >0.3 and minor allele frequency (MAF) >1%. We removed the effect of sex and age on batch effect-corrected metabolites levels, and used inverse normal transformation to normalize the residuals. We used RvTests (v20171009)(29) to test the association between genotype dosage and the various traits. We performed genetic association testing with bilirubin, retinopathy, aseptic necrosis, leg ulcers, survival status, painful crises, cholecystectomy, and eGFR in the CSSCD. Logistic regression model correcting for age, sex, SCD genotypes, HU usage and the first 10 principal components was employed to evaluate association between genotypes and binary traits. We used linear regression to test genetic associations with inverse normal-transformed eGFR, correcting for SCD genotypes, HU usage and the first 10 principal components.

### Mendelian Randomization

#### Instrument identification

Because of our reduced sample size, we selected instruments for MR analyses from large published mGWAS carried out in healthy individual of European ancestry. We identified metabolite-associated variants from the published meta-analysis of KORA and TwinsUK (N=6,056+1,768, 529 metabolites), as well as the whole-genome sequence metabolite association study in TwinsUK (N=1,960, 644 metabolites)(30, 31). We focused on these publications because they are the two largest published mGWAS to date. We selected subgenome-wide significant mGWAS SNPs (*P*<1×10^-5^) in order to maximize the phenotypic variance explained, and tested two MR models. The first MR model included all sub-genomewide significant SNPs as valid instruments. For the second MR model, we removed pleiotropic SNPs from the first model if they were associated with other metabolites at a Bonferroni-corrected *P*<0.05 threshold when considering the number of SNPs in model 1. Pleiotropic SNPs were identified by querying Phenoscanner (32).

#### Instrument pruning

We employed *PLINK1.9v5.2(33)* to identify independent SNP within 5-Mb window and linkage disequilibrium (LD) *r*^2^ <0.01 in the CSSCD. This provided us with a list of pseudo-independent variants.

#### Analysis

We used a two-sample MR approach to test the causal link between metabolites and SCD-related phenotypes. As described above, instruments and their effect sizes were selected from large European-ancestry mGWAS. We retrieved association results (effect sizes, standard errors) between instruments and SCD-related phenotypes from the large and clinically well-characterized CSSCD. All MR analyses were performed in R version 3.5.1 with the TwoSampleMR package (v0.4.22)(34). We used a multiplicative random-effect inverse variance-weighted (IVW) method in each MR analysis. For the analysis of L-glutamine and painful crises, we tested 2 models and defined statistical significance using a Bonferroni-corrected threshold of α≤0.025. All other analyses were exploratory and statistical significance was set at nominal α≤0.05. Additionally, we computed the weighted median (35), which selects the median MR estimate as the causal estimate, and MR-Egger (36), which allows the intercept to vary freely and therefore estimates the amount of horizontal pleiotropy, for all the analyses. Moreover, we utilized MR-PRESSO (Pleiotropy Residual Sum and Outlier)(37) to estimate the presence of horizontal pleiotropic bias and to calculate causal estimate adjusted for outliers for all reported results. Finally, we assess the validity of our significant results by conducting additional tests for horizontal pleiotropy, including Cochran’s Q statistic, MR-Egger intercept test of deviation from the null, and I^2^ heterogeneity statistic (13). Results from all MR analyses are available online at: http://www.mhi-humangenetics.org/dataset/MR_Analysis_SCD_everything.html.

### Genetic risk scores (GRS)

Using PLINK1.9v5.2 (33), we calculated the genetic risk scores for L-glutamine and 3-ureidopropionate in CSSCD, GEN-MOD and OMG. Effect size estimates from the two large mGWAS referenced in the MR analysis served as weights. Employing logistic regression, inverse-normal transformed GRS were associated to painful crisis adjusting for age, sex, the first 10 principal components (PCs), and SCD genotypes whenever appropriate. Employing linear regression, inverse-normal transformed GRS were associated to eGFR adjusting for the first 10 PCs and SCD genotypes whenever appropriate. Finally, in OMG and GEN-MOD, using linear regression, inverse-normal transformed GRS for L-glutamine and 3-ureidopropionate were associated to L-glutamine and 3-ureidopropionate metabolite levels, respectively. Both models were adjusted for age, sex, the first 10 PCs and SCD genotypes whenever appropriate.

## RESULTS

### Plasma metabolites in SCD patients

To identify metabolites that may be useful to predict or treat SCD complications, we measured plasma values of 129 known metabolites in 705 patients from the GEN-MOD and OMG cohorts (**Figure 1** and **Table 1**). Although our metabolomic experiment was performed at the same center, it was run in three batches so we applied stringent quality-control and batcheffect correction filters to avoid confounding (**Methods** and **Supplementary Figure 1**). The two main classes of metabolites that we measured were amino acids (33%) and lipids (30%), although we also captured carbohydrates, co-factors/vitamins, nucleotides, and energy-related metabolites (**Supplementary Figure 2** and **Supplementary Table 1**).

### Mendelian randomization supports a causal link between L-glutamine and SCD painful crises

L-glutamine therapy in SCD was previously shown to improve the NAD redox ratio, although this effect was not detected in a recent clinical trial (5, 38). Because we measured L-glutamine as part of our metabolomic experiment, we were interested to test association between its plasma levels and SCD-related complications or other clinically-relevant parameters. In GEN-MOD and OMG, we found no evidence of association between plasma L-glutamine levels and SCD complications, including painful crises (**Table 2**). However, L-glutamine levels were nominally associated with several hematological traits measured at baseline, including reduced hemoglobin concentration and RBC count (**Table 2**). For SCD complications, interpretation of these results is challenging because clinical events occurred before L-glutamine was measured, and this onetime metabolomic measure may not reflect life-long endogenous exposure to L-glutamine. For these reasons, we sought to further test the relationship between L-glutamine and SCD painful crises using MR.

**Table 2.** Associations between L-glutamine plasma levels and sickle cell disease (SCD)-related complications and other clinically relevant phenotypes. In participants from the GEN-MOD and OMG cohorts, we tested the association between L-glutamine levels measured in plasma and SCD-related complications or clinically relevant blood-based biomarkers. Dichotomous phenotypes were analyzed using logistic regression while correcting for age, sex, hydroxyurea (HU) usage, SCD genotypes and cohort affiliation. Quantitative phenotypes were corrected for age, sex, HU usage, SCD genotypes and cohort affiliation. They were inverse normal-transformed before being tested for association using linear regression. Odds ratio and effect sizes (Beta) are given per standard deviation change in L-glutamine plasma levels. LDH, lactate dehydrogenase; RBC, red blood cell; MCV, mean corpuscular volume; MCH, mean corpuscular hemoglobin; eGFR, estimated glomerular filtration rate; LDH, lactate dehydrogenase; CI, confidence interval; SE, standard error.

| **Complications** | **N** | **Odds ratio** | **95% CI** | **P-value** |
| --- | --- | --- | --- | --- |
| Painful crises | 619 | 1.06 | (0.90-1.24) | 0.52 |
| Survival | 529 | 1.01 | (0.75-1.35) | 0.79 |
| Aseptic necrosis | 617 | 0.97 | (0.97-1.16) | 0.76 |
| Cholecystectomy | 651 | 1.06 | (0.90-1.25) | 0.45 |
| Leg ulcer | 623 | 1.09 | (0.88-1.35) | 0.44 |
| Priapism | 448 | 1.11 | (0.88-1.4) | 0.39 |
| Retinopathy | 524 | 0.99 | (0.82-1.18) | 0.88 |
| **Renal Parameter** | **N** | **Beta** | **SE** | **P-value** |
| eGFR | 702 | -0.067 | 0.036 | 0.067 |
| **Blood Parameter** | **N** | **Beta** | **SE** | **P-value** |
| Bilirubin | 585 | 0.10 | 0.041 | 0.010 |
| Hematocrit | 697 | -0.08 | 0.035 | 0.019 |
| Hemoglobin | 685 | -0.098 | 0.035 | 0.0048 |
| LDH | 579 | 0.078 | 0.039 | 0.044 |
| MCH | 626 | 0.067 | 0.035 | 0.053 |
| MCV | 697 | 0.07 | 0.032 | 0.03 |
| RBC | 698 | -0.11 | 0.033 | $7.1 \times 10^{-4}$ |

**Table 3.** Nominally significant associations between survival and plasma levels of 129 metabolites in sickle cell disease (SCD). In participants from the GEN-MOD and OMG cohorts, we tested the association between metabolite levels measured in plasma and prospectively ascertained survival (529 SCD patients: 54 deaths). We analyzed the data using logistic regression while correcting for age, sex, hydroxyurea (HU) usage, SCD genotypes and cohort affiliation. Metabolite levels were inverse normal-transformed so that odds ratios are given per standard deviation change in plasma metabolite levels. Only quinolinic acid remains significant after permutations to account for the number of tests performed. OR, odds ratio; CI, confidence interval.

| Metabolite | OR | 95% CI | P-value | Permuted P-value | Permuted P-value after adjusting for eGFR |
| --- | --- | --- | --- | --- | --- |
| Quinolinic acid | 1.76 | (1.29 - 2.41) | $3.8 \times 10^{-4}$ | 0.03 | 0.89 |
| Cytosine | 1.60 | (1.19 - 2.15) | $1.8 \times 10^{-3}$ | 0.15 | 0.16 |
| Indoxyl sulfate | 1.59 | (1.17 - 2.17) | $3.1 \times 10^{-3}$ | 0.24 | 1.0 |
| 2-trans,4-cis-Decadienoylcarnitine | 1.53 | (1.14 - 2.05) | $4.2 \times 10^{-3}$ | 0.3 | 1.0 |
| Cytidine | 1.52 | (1.14 - 2.04) | $5.0 \times 10^{-3}$ | 0.34 | 0.97 |
| Asymmetric dimethylarginine | 1.53 | (1.13 - 2.07) | $5.6 \times 10^{-3}$ | 0.38 | 0.78 |
| L-Cystathionine | 1.56 | (1.13 - 2.14) | $6.6 \times 10^{-3}$ | 0.42 | 1.0 |
| Alpha-Lactose | 1.47 | (1.1 - 1.97) | $9.2 \times 10^{-3}$ | 0.53 | 1.0 |
| D-Glucuronic acid | 1.5 | (1.11 - 2.05) | $9.3 \times 10^{-3}$ | 0.54 | 0.97 |
| Ureidopropionic acid | 1.47 | (1.1 - 1.97) | $1.0 \times 10^{-2}$ | 0.56 | 1.0 |

Instrument strength plays a critical role in the validity of MR analyses. Although we measured L-glutamine levels in 705 SCD patients, we wanted to take advantage of existing and well-powered mGWAS for the selection of the best metabolite-associated SNPs to use as MR instruments (30, 31). However, these mGWAS were carried out in Europeans, whereas SCD patients in our cohorts are of African-descent, raising the question whether we could use SNPs found in Europeans as MR instruments for phenotypes observed in African-ancestry SCD patients. To validate this strategy, we tested the well-known causal link between bilirubin levels in serum and gallstones leading to surgical removal of the gallbladder (cholecystectomy), a complication often observed in SCD patients (39). From a GWAS of serum bilirubin levels in 9,464 individuals of European-ancestry, we selected 10 SNPs as MR instruments (40). In the large and well-characterized CSSCD (**Table 1**), we tested the association between these SNPs and bilirubin levels or cholecystectomy, and replicated the strong association between these phenotypes and the *UGT1A1* locus (**Supplementary Table 2**). The two-sample inverse variance-weighted (IVW) MR analysis confirmed that high bilirubin levels causes gallbladder disease in SCD: a one standard deviation increase in genetically-controlled bilirubin levels was associated with a 6-fold increase in the risk of cholecystectomy in the CSSCD (odds ratio (OR) [95% confidence interval] = 6.0 [2.8-17.0], *P*_IVW_=1.9×10^-6^)(**Supplementary Table 3**).

From the available mGWAS results (30, 31), we identified 51 SNPs associated with plasma L-glutamine levels at *P*<5×10^-5^ that were available in the CSSCD genetic dataset. Single variant and polygenic trait score association results are available in **Supplementary Table 4** (25). Using these 51 SNPs as instruments in a two-sample IVW MR analysis, we did not detect a causal association between L-glutamine and painful crises (Model 1: OR = 0.81 [0.63-1.00], *P*_IVW_=0.086)(**Figure 2**). When we excluded 24 pleiotropic SNPs (**Methods**) and repeated the analysis with the remaining 27 SNPs, the MR association with painful crises was significant: a one standard deviation increase in genetically-controlled L-glutamine levels was associated with a 32% reduction in the risk of painful crises in the CSSCD (Model 2: OR=0.68 [0.52-0.89], *P*_IVW_=0.0048)(**Figure 2**). MR analyses using the sensitivity tests MR-Egger and weighted-median did not yield significant associations for Model 2, suggesting insufficient statistical power for these tests (13) or potential residual pleiotropy (**Supplementary Table 5**). We repeated the MR analysis in the GEN-MOD and OMG cohorts: although the direction of the effect of the GEN-MOD+OMG meta-analysis indicated a protective effect of L-glutamine on painful crises (OR=0.82 [0.54-1.34]), the result was not significant (*P*_IVW_=0.54), presumably due to limited power given the smaller sample size (**Supplementary Table 5**). In secondary MR analyses, we found no evidence of causal associations between L-glutamine SNPs and several other SCD complications (**Supplementary Table 5**).

**Figure 2.**
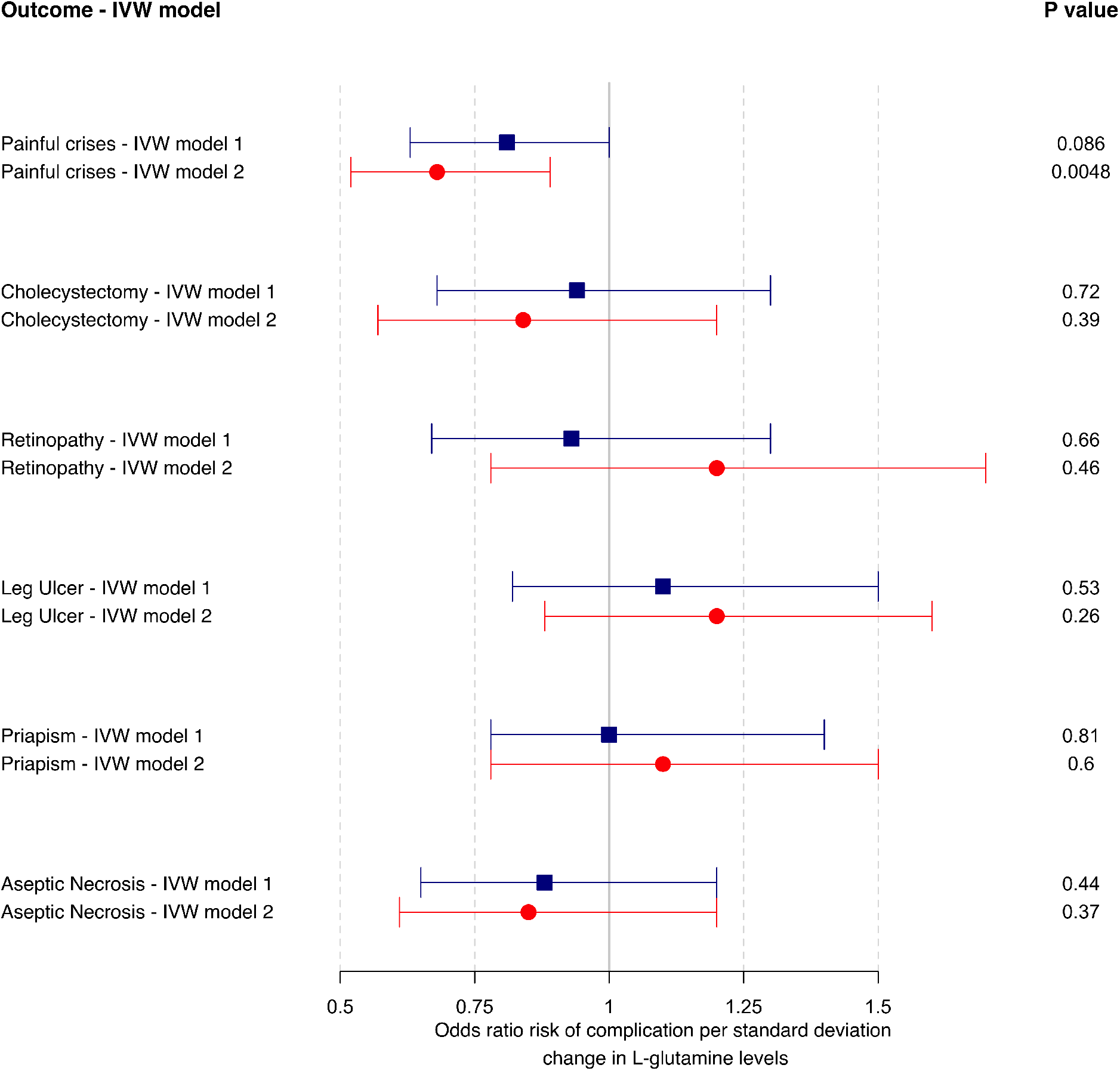
Mendelian randomization (MR) analysis of L-glutamine with sickle cell disease (SCD) painful crises. Forest plot of MR evaluating the causal relationship between L-glutamine levels and painful crises in SCD patients. Effect sizes and standard errors of 51 variants associated with L-glutamine were retrieved from large European mGWAS. Associations statistics between these 51 variants and SCD complications were calculated in the large prospective and well-characterized CSSCD. In model 1, we considered all 51 SNPs as instruments, whereas model 2 only included 27 variants not associated with other metabolites (Methods). The MR effect size estimates and 95% confidence intervals were calculated using the inverse variance-weighted (IVW) random effect method.

### Potential causal link between 3-ureidopropionate and kidney function in SCD

We tested 6 SCD-related complications as well as eGFR against the levels of the 129 known metabolites measured in GEN-MOD and OMG. In total, we found 65 metabolites with *P*_perm_ ≤0.05, including 61 metabolites associated with eGFR (**Figure 3** and **Supplementary Table 6**). There was a strong association between eGFR and creatinine levels, which serves as an internal control given that we use this metabolite to calculate eGFR. Most of these metabolites have never been linked to SCD, and may therefore represent potential novel biomarkers of disease severity.

**Figure 3.**
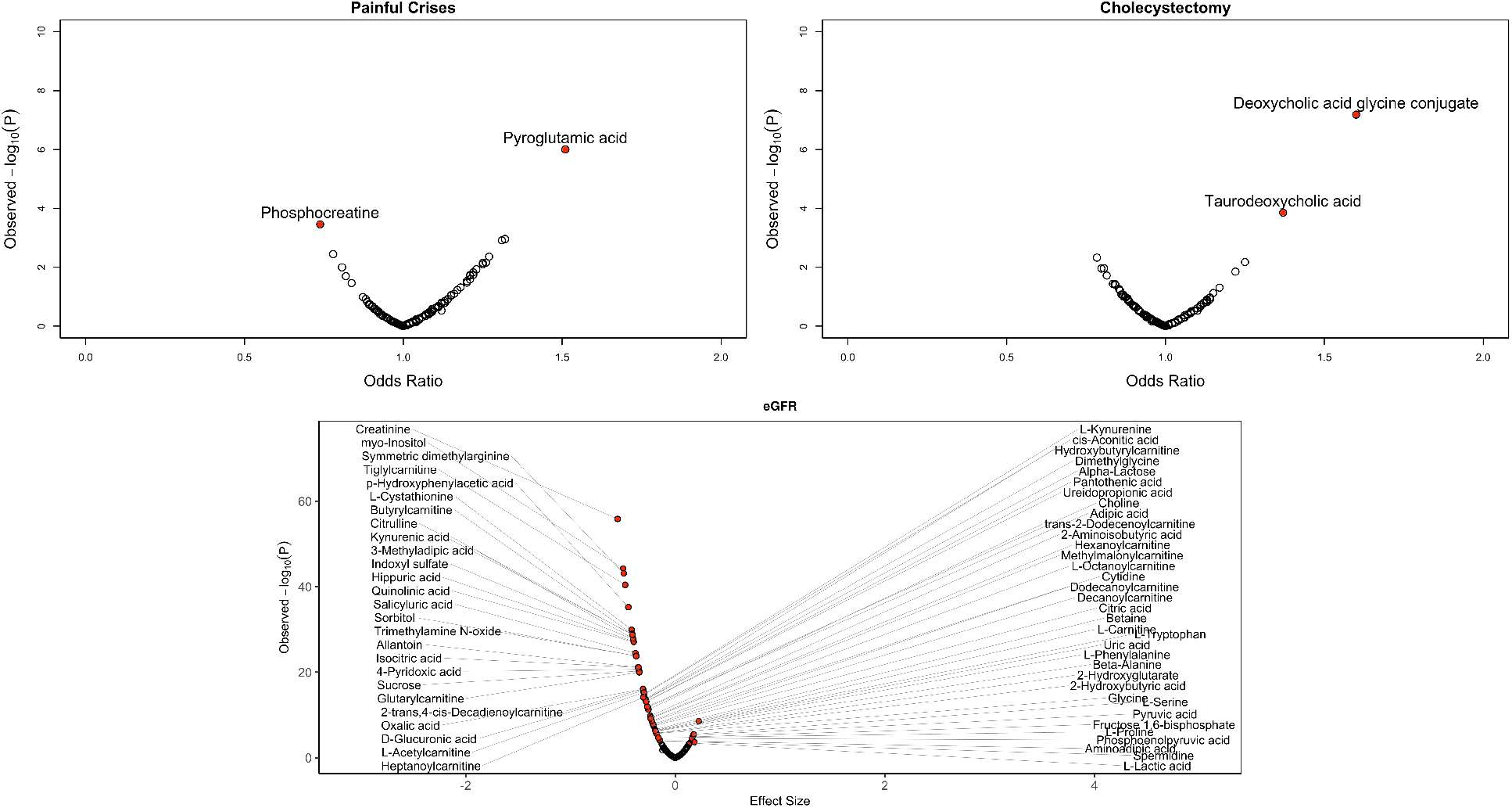
Known metabolites associated with SCD complications and estimated glomerular filtration rate (eGFR). We tested 129 metabolites against clinical complications by logistic regression (linear regression for quantitative eGFR). On the *x*-axis, we report odd ratios (effect sizes for eGFR) in metabolite standard deviation units, whereas the, *y*-axis presents the observed analytical P-values. Red circles highlight metabolites with *P*_perm_ <0.05 calculated using 100,000 permutations. In total, we found 2 metabolites for painful crises, 2 metabolite with cholecystectomy, and 61 metabolites for eGFR.

Using the same strategy as for L-glutamine, we derived MR instruments for 48 of the 66 metabolites identified in the pairwise analyses with SCD phenotypes; there were no significant mGWAS variants for the remaining 18 metabolites. Across these 48 tests, we identified a single nominally significant association in our two-sample MR analyses involving eGFR and 3-ureidoproprionate levels (see URL for all available MR results, including sensitivity tests). In a European mGWAS (31), we retrieved 22 SNPs associated with 3-ureidoproprionate levels, including 16 that were not pleiotropic (**Supplementary Table 7**). Our results indicate that a one standard deviation increase in genetically-controlled 3-ureidopropionate levels was associated with improved eGFR of 0.07 mL/min per 1.172 m^2^ (*P*_IVW-model1_=8.7×10^-4^; *P*_IVW-model2_=9.7×10^-4^) (**Figure 4** and **Supplementary Tables 8**). The sensitivity analyses did not allow us to exclude the possibility of confounding due to pleiotropy (**Figure 4** and **Supplementary Tables 8**). Furthermore, we could not replicate the MR result in GEN-MOD and OMG, indicating that larger SCD cohorts are needed to confirm the causal link between 3-ureidoproprionate and eGFR (**Supplementary Table 7**).

**Figure 4.**
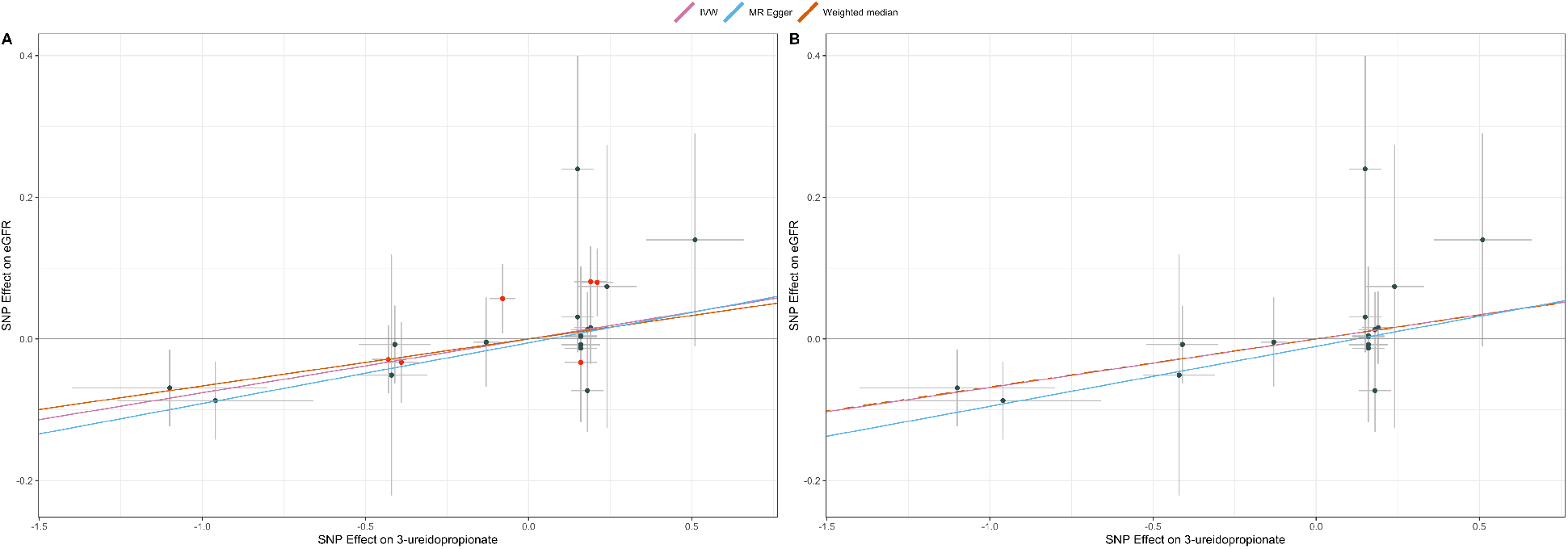
3-ureidopropionate causally influences estimated glomerular filtration rate (eGFR) in sickle cell disease (SCD) patients. (**A**) Mendelian randomization (MR) plot comparing the effects of SNPs on 3-ureidopropionate in Europeans (*x*-axis) and eGFR in SCD patients (*y*-axis). The slope of each line corresponds to the MR effect for each method (inverse variance-weighted (IVW), MR-Egger or weighted median). Data are expressed as effect sizes with 95% confidence intervals. SNPs in red are pleiotropic. (**B**) Same as **A**, except that we removed pleiotropic variants.

### Predicting survival status using baseline metabolite levels

Given the clinical heterogeneity that characterizes this disease, being able to predict which SCD patients will follow a severe clinical course could be extremely useful. Thus, we decided to explore the prognostic value of plasma metabolites in SCD. As discussed above, the data currently available in GEN-MOD and OMG are largely retrospective. However, we could prospectively ascertain SCD severity using a simple definition based on survival status during the follow-up period (Table 1 and Methods). We identified 10 metabolites that were nominally associated with survival status, but only quinolinic acid remained significant after permutations to account for the number of tests performed. For all 10 metabolites, increased levels were associated with increased risk of death, and for all but cytosine levels, the effect on survival was mediated by an association with eGFR. Quinolinic acid is a product of the kynurenine pathway, which also metabolizes the amino acid tryptophan.

## DISCUSSION

While the cause of SCD has been known for over a century, treatment options are limited and it is extremely difficult to predict which patients will have a more severe presentation of the disease. To continue to address these challenges, we performed the largest metabolomic study in SCD patients, measuring 129 known metabolites in 705 participants. Our effort was motivated by recent successes using this approach to find new prognostic biomarkers and potential drug targets for human diseases (41). By combining metabolite profiles with mGWAS results, we could use MR methods to test causality between metabolites and SCD-related complications. Although a few metabolites have previously been implicated in SCD clinical heterogeneity (adenosine, S1P, 2,3-BPG)(19–21), we did not measure them and could therefore not replicate their associations in our dataset. However, we identified a promising causal relationship between L-glutamine levels and painful crises, consistent with recent results from a phase 3 clinical trial(5).

Our analyses also highlighted 3-ureidoproprionate, an intermediate in the metabolism of uracil, as a potential positive modulator of eGFR. Interpretation of this result is difficult because little is known about this metabolite and the result was not replicated in additional SCD patients. Mutations in the gene *UPB1*, which encodes the enzyme that transforms 3-ureidopropionate into beta-alanine, cause beta-ureidopropionase deficiency, a rare monogenic disease characterized by high plasma levels of 3-ureidoproprionate (42). Only a few patients with this disease have so far been described and they presented mostly with neurologic development issues. However, there is no report of abnormal glomerular filtration rate or other kidney defects in these patients. We propose that future MR replication in independent SCD cohorts and animal studies could be extremely useful to investigate the possible role of 3-ureidoproprionate in regulating kidney functions, and in particular whether raising 3-ureidopropionate levels could improve glomerular filtration rate in SCD patients.

Our study presents with a few limitations. First, our statistical power to detect heterogeneity (for instance due to horizontal pleiotropy in our MR analyses) and to replicate our main findings was limited because there are few large, well-characterized and genotyped SCD cohorts available. Second, we measured metabolite levels in plasma, but their levels in RBC could have provided complementary information (in particular for L-glutamine). Third, we used MR instruments derived from mGWAS performed in Europeans to test for causality in African-ancestry SCD patients. There have been many reports on the transferability (or lack thereof) of GWAS findings across ancestries (43). We used the well-known relationship between bilirubin levels and gallbladder disease to show that our approach can work. However, it is likely that having access to large mGWAS results in African-ancestry populations would provide better instruments, and may lead to the identification of additional causal link between metabolites and SCD phenotypes by MR.

One characteristic of our study is that we measured metabolites in SCD cohorts that have mostly collected retrospective clinical data. One exception is information on the survival status of the participants. Using a simple linear model, we found a significant association between prospectively-ascertained survival status and baseline quinolinic acid levels. This association was mediated by eGFR, consistent with our previous observation that quinolinic acid levels correlate with rapid renal function decline in SCD patients (24). In the future, it will be interesting and important to test whether metabolites predict other complications in prospective SCD cohorts. In conclusion, our results motivate future experiments to integrate metabolite profiles and other orthogonal omics datasets (*e.g*. genetics) to build better predictors of SCD-related complications and overall severity.

## Author contributions

A.E.A.-K., M.J.T and G.L. conceived this study. C.B.C. performed plasma metabolites profiling. Y.I. performed Mendelian randomization analyses. Y.I. and M.G. performed genetic analyses and genetic risk association analyses. Y.I. performed statistical analyses. G.L. supervised this work. Y.I. and G.L. wrote the manuscript with input from all authors.

## Acknowledgments

We thank all participants for their contribution to this project. We also thank Adil Harroud and Brent Richards for advices on the Mendelian randomization analyses. G.L. is funded by the Canadian Institutes of Health Research (CIHR, PJT #156248), the Doris Duke Charitable Foundation, and the Canada Research Chair program. GEN-MOD sample and data collection were supported by NIH grant HL-68922. A.A-K. M.J.T. and establishment and analysis of the OMG cohort has been funded by NHLBI (R01HL68959, HL79915, HL70769, HL87681).

## URL

All Mendelian randomization results are available at: http://www.mhi-humangenetics.org/dataset/MR_Analysis_SCD_everything.html

