## Supplemental Material Text for "Causal role of L-glutamine in sickle cell disease painful crises: a Mendelian randomization analysis"

**Affiliations**
[1]Montreal Heart Institute, Montréal, Québec, Canada.
[2]Faculté de Médecine, Université de Montréal, Montréal, Québec, Canada
[3]Duke Molecular Physiology Institute, Duke University Medical Center, Durham, North Carolina, USA
[4]Red Cell Genetic Disease Unit, Hôpital Henri-Mondor, Assistance Publique-Hôpitaux de Paris (AP-HP), Université Paris Est, IMRB - U955 - Equipe no 2, Créteil, France
[5]Department of Laboratory Medicine, Boston Children's Hospital, Boston, Massachusetts
[6]Broad Institute, Cambridge, MA, USA
[7]Boston Children's Hospital, Boston, MA, USA
[8]Department of Medicine, Division of Hematology, Duke University Medical Center, Durham, North Carolina, USA

**Corresponding author**
Guillaume Lettre
Montreal Heart Institute
5000 Belanger Street
Montreal, Quebec, Canada
.

**Supplementary Table 1. List of 129 known metabolites measured in the plasma of sickle cell disease (SCD) patients.** For each metabolite, we provide the pathway, super pathway and identifier from the human metabolome database (HMDB ID), as well as mean and standard deviation in the GEN-MOD and OMG SCD cohorts.

SEE EXCEL FILE

**Supplementary Table 2. Genetic associations for bilirubin levels and cholecystectomy.** SNPs (effect alleles) are the lead variants in each gene region identified in a GWAS for bilirubin levels in European-ancestry individuals[1]. Association effect sizes with bilirubin levels in the CSSCD are in standard deviation units, and association effect sizes with bilirubin levels in Johnson et *al*. are in log-transformed bilirubin units. Association effect size with cholecystectomy in the CSSCD are log odds ratios from logistic regression analyses. These variants were selected because of their strong association ($P<2.0 \times 10^{-5}$) with bilirubin levels in Johnson et *al*. *These variants were identified in the Johnson et *al*. study, but because their minor allele frequency (MAF) <1% they were excluded from the Mendelian randomization analysis.

| | | Association result with bilirubin-associated variants (effect allele frequency, effect size (β), P-value, sample size) | | |
|---|---|---|---|---|
| **SNP** | **Gene region** | **Bilirubin (Johnson et al)** | **Bilirubin (CSSCD)** | **Cholecystectomy Risk (CSSCD)** |
| rs6742078(T) | UGT1A1 | (0.31, 0.23, P = 5e-324, N = 8988) | (0.44, 0.43, P = 9.7e-21, N = 930) | (0.43, 0.44, P = 0.00091, N = 1084) |
| rs12714207(T) | KRCC1 | (0.33, -0.033, P = 5.3e-07, N = 8988) | (0.52, 0.047, P = 0.31, N = 930) | (0.53 -0.065, P = 0.63, N = 1084) |
| rs1986655(A) | intergenic | (0.15, -0.035, P = 2e-06, N = 8988) | (.96, 0.026, P = 0.82, N = 930) | (0.96, 0.33, P = 0.33, N = 1084) |
| rs12206204(T)/ rs113892814(A)* | histone cluster | (0.015, 0.15, P = 7.5e-07, N = 8988) | (0.0022, -0.27, P = 0.59, N = 930) | (0.00046/0.0028*, 1.8, P = 0.28, N = 1084) |
| rs9380833(T) | KCNK5 | (0.027, 0.079, P = 1.6e-05, N = 8988) | (0.11, -0.022, P = 0.76, N = 930) | (0.11, 0.12, P = 0.55, N = 1084) |
| rs4236644(A) | SEMA3C | (0.27, -0.031, P = 2.1e-06, N = 8988) | (0.52, -0.065, P = 0.15, N = 930) | (0.51, 0.039, P = 0.77, N = 1084) |
| rs12337836(A)* | PRG-3, BAAT | (0.076, 0.053, P = 1.3e-05, N = 8988) | (0.0054, -0.14, P = 0.66, N = 930) | (0.0055, 0.03, P = 0.98, N = 1084) |
| rs16928809(A)* | SLC22A18 | (0.096, 0.051, P = 1.1e-07, N = 8988) | (0.0038, 0.11, P = 0.77, N = 930) | (0.0037, 0.99, P = 0.35, N = 1084) |
| rs4149056(T) | SLCO1B1 | (0.15, -0.053, P = 6.7e-13, N = 8988) | (0.978, -0.26, P = 0.11, N = 930) | (0.98, -0.23, P = 0.63, N = 1084) |
| rs4773330(A) | ARHGEF7 | (0.12, -0.036, P = 7.7e-06, N = 8988) | (0.17, 0.03, P = 0.63, N = 930) | (0.17, 0.15, P = 0.41, N = 1084) |
| rs7173819(A) | intergenic | (0.12, 0.035, P = 1.2e-05, N = 8988) | (0.79, -0.021, P = 0.71, N = 930) | (0.79, 0.19, P = 0.26, N = 1084) |
| rs12923103(A) | intergenic | (0.32, 0.027, P = 1.4e-05, N = 8988) | (0.17, 0.037, P = 0.55, N = 930) | (0.17, 0.12, P = 0.49, N = 1084) |
| rs4410172 (C) | BC051727 | (0.24, 0.026, P = 1.9e-05, N = 8988) | (0.097, -0.11, P = 0.16, N = 930) | (0.098, 0.21, P = 0.37, N = 1084) |

**Supplementary Table 3. Mendelian randomization (MR) results of bilirubin levels with cholecystectomy.** As instruments for our MR analyses, we used SNPs identified by Johnson et *al*. in a genome-wide association study (GWAS) for bilirubin levels in 9,464 individuals of European ancestry[1]. From the list of 15 bilirubin-associated variants from the GWAS, we kept 10 SNPs that were in linkage equilibrium (rs6742078, rs12714207, rs1986655, rs9380833, rs4236644, rs4149056, rs4773330, rs7173819, rs12923103, rs4410172). We analyzed cholecystectomy in 1,294 participants from the CSSCD. We performed a two-sample MR analysis using exposure effect sizes from the bilirubin GWAS of Johnson et *al*. (GWAS_beta). For the two-sample MR analysis, effect sizes of the MR are odds to develop cholecystectomy per 27% increase in bilirubin levels.

| Method | Number of variants | Odds ratio (95% confidence interval) and P-value |
|---|---|---|
| **Inverse variance-weighted (IVW)** | 10 | 6.0 (2.8-17)<br>$P=1.9 \times 10^{-6}$ |
| **MR-Egger** | 10 | 7.0 (1.7-29)<br>P=0.027 |
| **Weighted median** | 10 | 6.4 (2.1-20)<br>$P=1.2 \times 10^{-3}$ |

**Supplementary Table 4. Genetic associations between 51 L-glutamine-associated SNPs and painful crises.** SNPs (effect alleles) are the lead variants in each gene region identified in mGWAS for L-glutamine levels in Europeans (*P* $<5.0\text{x}10^{-5}$)[2,3]. Association effect sizes with painful crises in the CSSCD and GEN-MOD are log odds ratios from logistic regression. *These variants were identified to be pleiotropic by Phenoscanner queries and were excluded from model 2 (**Methods**). At the bottom of the table, we also provide association results between polygenic trait scores (PTS) calculated using 51 L-glutamine-associated SNPs ($PTS_{51SNPs}$) or after excluding pleiotropic variants ($PTS_{27SNPs}$). For the PTS, the effect size is per PTS standard deviation units.

| | | **Association result with L-glutamine-associated variants (effect allele frequency, effect size (beta or OR), P-value, sample size** | | | |
|---|---|---|---|---|---|
| **SNP (hg19)** | **Gene region** | **L-glutamine (Shin et al/Long et al)** | **L-glutamine (Meta-analysis GEN-MOD-OMG)** | **Painful crises (CSSCD)** | **Painful crises (Meta-analysis GEN-MOD-OMG)** |
| rs524219(G)/1:100821493 | CDC14A | (0.93, -0.28, P = 4.7e-06, N = 1958) | (0.02, -0.21, P= 0.33, N = 651) | (0.99 ,0.59, P = 0.24, N = 1101) | (0.98, -0.91, P = 0.22, N = 575) |
| rs11166473(T)/1:101010015* | GPR88 | (0.92, -0.36, P = 9.3e-08, N = 1958) | (0.48, 0.01, P= 0.85, N = 651) | (0.47 ,-0.12, P = 0.31, N = 1101) | (0.49, -0.11, P = 0.56, N = 575) |
| rs2811981(A)/1:23950147* | MDS2 | (0.89, -0.12, P = 3.8e-06, N = 1958) | (0.87, 0.09, P= 0.28, N = 651) | (0.88 ,-0.12, P = 0.49, N = 1101) | (0.87, -0.11, P = 0.69, N = 575) |
| rs3127550(A)/1:49431909* | AGBL4 | (0.49, 0.19, P = 5.7e-06, N = 1958) | (0.18, -0.02, P= 0.77, N = 651) | (0.17 ,-0.021, P = 0.89, N = 1101) | (0.19, -0.52, P = 0.03, N = 575) |
| rs7394051(T)/10:122008026* | intergenic | (0.7, 0.0055, P = 2.2e-06, N = 7372) | (0.72, 0.06, P= 0.39, N = 651) | (0.76 ,-0.052, P = 0.7, N = 1101) | (0.72, 0.14, P = 0.51, N = 575) |
| rs10762121(T)/10:68708291 | CTNNA3 | (0.18, -0.0094, P = 3.1e-05, N = 1768) | (0.44, 0.04, P= 0.45, N = 651) | (0.4 ,-0.018, P = 0.88, N = 1101) | (0.44, 0.04, P = 0.84, N = 575) |
| rs11596604(A)/10:87141944* | intergenic | (0.024, 0.032, P = 2.5e-06, N = 1768) | (0, -0.43, P= 0.46, N = 401) | (0.01 ,0.16, P = 0.78, N = 1101) | (0, -2.5, P = 0.07, N = 325) |
| rs7078003(T)/10:99359412* | HOGA1 | (0.17, 0.0087, P = 1.8e-06, N = 7372) | (0.11, -0.01, P= 0.93, N = 651) | (0.12 ,-0.12, P = 0.5, N = 1101) | (0.12, -0.21, P = 0.48, N = 575) |
| rs7131407(T)/11:128431418 | ETS1 | (0.13, 0.0082, P = 3.2e-05, N = 7372) | (0.16, 0.02, P= 0.82, N = 651) | (0.16 ,0.29, P = 0.074, N = 1101) | (0.16, 0.19, P = 0.46, N = 575) |
| rs10431159(A)/11:70967219 | SHANK2 | (0.65, -0.0047, P = 4.8e-05, N = 7372) | (0.1, 0.09, P= 0.29, N = 651) | (0.15 ,-0.061, P = 0.71, N = 1101) | (0.1, -0.06, P = 0.86, N = 575) |
| rs17666239(A)/12:47194757* | SLC38A4 | (0.078, 0.0092, P = 1.7e-05, N = 7372) | (0.03, 0.1, P= 0.53, N = 651) | (0.035 ,0.24, P = 0.45, N = 1101) | (0.03, -0.69, P = 0.22, N = 575) |
| rs735246(A)/12:52547743 | intergenic | (0.83, 0.0083, P = 4.8e-05, N = 7372) | (0.82, -0.05, P= 0.49, N = 651) | (0.81 ,-0.11, P = 0.42, N = 1101) | (0.82, -0.08, P = 0.74, N = 575) |
| rs774044(T)/12:56837979* | TIMELESS | (0.055, -0.015, P = 5.6e-07, N = 7372) | (0.05, -0.11, P= 0.4, N = 651) | (0.045 ,0.093, P = 0.74, N = 1101) | (0.05, -0.17, P = 0.7, N = 575) |
| rs7313455(A)/12:56853231* | MIP | (0.44, 0.0062, P = 2.1e-08, N = 7372) | (0.1, 0.11, P= 0.29, N = 651) | (0.13 ,-0.026, P = 0.88, N = 1101) | (0.06, 0.2, P = 0.63, N = 575) |

| | | | | | |
|---|---|---|---|---|---|
| rs2657879(A)/12:56865338* | GLS2 | (0.82, 0.015, P = 6.1e-18, N = 7372) | (0.96, 0.19, P= 0.15, N = 651) | (0.94 ,0.25, P = 0.32, N = 1101) | (0.96, -0.4, P = 0.41, N = 575) |
| rs12232026(A)/12:56960766* | RBMS2 | (0.12, -0.0097, P = 4.4e-06, N = 7372) | (0.17, 0.12, P= 0.1, N = 651) | (0.14 ,-0.32, P = 0.049, N = 1101) | (0.16, 0.12, P = 0.64, N = 575) |
| rs941893(T)/14:100542061* | EVL | (0.74, -0.0049, P = 4.1e-05, N = 7372) | (0.26, -0.01, P= 0.86, N = 651) | (0.27 ,0.013, P = 0.92, N = 1101) | (0.25, -0.08, P = 0.72, N = 575) |
| rs144325715(A)/14:95948631 | intergenic | (0.021, -0.74, P = 5.3e-06, N = 1958) | (0.04, -0.12, P= 0.46, N = 651) | (0.044 ,0.25, P = 0.4, N = 1101) | (0.04, 0.42, P = 0.38, N = 575) |
| rs11636988(A)/15:26822814* | GABRB3 | (0.57, -0.16, P = 6.5e-06, N = 1958) | (0.08, 0.14, P= 0.18, N = 651) | (0.13 ,-0.0023, P = 0.99, N = 1101) | (0.07, -0.25, P = 0.53, N = 575) |
| rs1910151(A)/15:38158699* | NA | (0.64, 0.0048, P = 2.2e-05, N = 7372) | (0.64, -0.05, P= 0.42, N = 651) | (0.64 ,0.11, P = 0.36, N = 1101) | (0.64, 0.34, P = 0.08, N = 575) |
| rs35150605(TAAC)/15:88047276 | RP11-648K4.2 | (0.25, -0.15, P = 6.3e-06, N = 1958) | (0.7, -0.03, P= 0.58, N = 651) | (0.3 ,0.037, P = 0.77, N = 1101) | (0.28, 0.09, P = 0.66, N = 575) |
| rs2560409(C)/16:24099496 | PRKCB | (0.51, 0.26, P = 1.4e-06, N = 1958) | (0.84, 0.19, P= 0.01, N = 651) | (0.18 ,-0.12, P = 0.43, N = 1101) | (0.14, -0.41, P = 0.13, N = 575) |
| rs16977047(T)/16:27924612* | GSG1L | (0.7, -0.0056, P = 1.1e-06, N = 7372) | (0.78, -0.01, P= 0.84, N = 651) | (0.77 ,-0.11, P = 0.43, N = 1101) | (0.8, -0.12, P = 0.61, N = 575) |
| rs12447776(T)/16:84053027 | SLC38A8 | (0.019, 0.029, P = 4.1e-05, N = 1768) | (0.05, -0.09, P= 0.49, N = 651) | (0.042 ,0.057, P = 0.85, N = 1101) | (0.06, -0.19, P = 0.65, N = 575) |
| rs9912445(A)/17:37202603 | LRRC37A11P | (0.79, 0.0052, P = 1.7e-05, N = 7372) | (0.71, 0.08, P= 0.17, N = 651) | (0.72 ,-0.17, P = 0.17, N = 1101) | (0.7, 0.3, P = 0.12, N = 575) |
| rs8069305(A)/17:53638670 | CTD-2033D24.2 | (0.68, -0.0047, P = 4.9e-05, N = 7372) | (0.14, 0.1, P= 0.18, N = 651) | (0.19 ,0.031, P = 0.84, N = 1101) | (0.16, 0.17, P = 0.49, N = 575) |
| rs4798682(G)/18:8738265 | SOGA2 | (0.74, 0.15, P = 5e-06, N = 1958) | (0.11, -0.15, P= 0.1, N = 651) | (0.87 ,-0.17, P = 0.33, N = 1101) | (0.89, 0.2, P = 0.52, N = 575) |
| rs73971292(C)/2:161821810 | intergenic | (0.023, -0.65, P = 5.1e-06, N = 1958) | (0.97, 0.13, P= 0.43, N = 651) | (0.025 ,0.15, P = 0.69, N = 1101) | (0.02, -0.68, P = 0.3, N = 575) |
| rs780093(T)/2:27742603* | GCKR | (0.4, -0.0058, P = 1.9e-07, N = 7372) | (0.12, -0.07, P= 0.42, N = 651) | (0.14 ,0.15, P = 0.37, N = 1101) | (0.12, 0.03, P = 0.93, N = 575) |
| rs2199619(T)/2:28854958* | PLB1 | (0.28, 0.0054, P = 5.2e-06, N = 7372) | (0.21, -0.11, P= 0.12, N = 651) | (0.22 ,-0.11, P = 0.45, N = 1101) | (0.19, -0.01, P = 0.97, N = 575) |
| rs992580(C)/20:15246472* | MACROD2 | (0.44, 0.15, P = 6.2e-06, N = 1958) | (0.53, 0.0036, P= 0.95, N = 651) | (0.47 ,-0.0083, P = 0.94, N = 1101) | (0.47, -0.12, P = 0.52, N = 575) |
| rs6137021(A)/20:20375560* | RALGAPA2 | (0.67, -0.0083, P = 4.5e-05, N = 1768) | (0.2, 0.17, P= 0.01, N = 651) | (0.23 ,-0.32, P = 0.023, N = 1101) | (0.18, 0.45, P = 0.05, N = 575) |
| rs2425059(C)/20:33912371* | UQCC1 | (0.38, 0.15, P = 5.6e-06, N = 1958) | (0.4, -0.01, P= 0.91, N = 651) | (0.56 ,-0.12, P = 0.31, N = 1101) | (0.63, -0.35, P = 0.08, N = 575) |
| rs2948828(A)/3:124811942 | SLC12A8 | (0.56, 0.0053, P = 2.2e-05, N = 7372) | (0.75, -0.0033, P= 0.96, N = 651) | (0.75 ,0.056, P = 0.67, N = 1101) | (0.76, -0.19, P = 0.39, N = 575) |
| rs73168973(T)/3:151691599 | intergenic | (0.21, -0.23, P = 4e-07, N = 1958) | (0.06, 0.03, P= 0.81, N = 651) | (0.079 ,-0.18, P = 0.42, N = 1101) | (0.06, -0.46, P = 0.28, N = 575) |
| rs4699183(A)/4:106444435 | AC004066.2 | (0.9, -0.27, P = 6.2e-06, N = 1958) | (0.93, 0.29, P= 0.0086, N = 651) | (0.92 ,0.082, P = 0.71, N = 1101) | (0.93, 0.51, P = 0.16, N = 575) |

| | | | | | |
|---|---|---|---|---|---|
| rs138354882(A)/4:171429817 | intergenic | (0.028, -0.36, P = 4.5e-06, N = 1958) | (0.1, -0.06, P= 0.52, N = 651) | (0.09 ,0.24, P = 0.25, N = 1101) | (0.1, 0.04, P = 0.91, N = 575) |
| rs7667615(C)/4:182921992* | AC108142.1 | (0.26, -0.17, P = 6.1e-06, N = 1958) | (0.81, -0.0035, P= 0.96, N = 651) | (0.17 ,-0.031, P = 0.84, N = 1101) | (0.18, 0.16, P = 0.52, N = 575) |
| rs542300(A)/6:12252237 | intergenic | (0.51, -0.13, P = 7.2e-06, N = 1958) | (0.54, 0.07, P= 0.22, N = 651) | (0.54 ,-0.064, P = 0.59, N = 1101) | (0.54, 0.32, P = 0.07, N = 575) |
| rs9478369(A)/6:153269698* | intergenic | (0.25, 0.0049, P = 4.1e-05, N = 7372) | (0.56, 0.0019, P= 0.97, N = 651) | (0.51 ,0.052, P = 0.66, N = 1101) | (0.57, -0.14, P = 0.44, N = 575) |
| rs71569656(C)/6:22903627 | RP1-209A6.1 | (0.28, -0.14, P = 8.9e-06, N = 1958) | (0.92, -0.01, P= 0.91, N = 651) | (0.095 ,0.42, P = 0.04, N = 1101) | (0.07, -0.46, P = 0.23, N = 575) |
| rs2748991(T)/6:52596516 | intergenic | (0.45, -0.007, P = 2.3e-06, N = 5604) | (0.33, 0.16, P= 0.0096, N = 651) | (0.34 ,-0.091, P = 0.45, N = 1101) | (0.33, -0.2, P = 0.34, N = 575) |
| rs1582256(C)/7:126634804 | GRM8 | (0.51, -0.18, P = 8.5e-06, N = 1958) | (0.5, -0.07, P= 0.19, N = 651) | (0.49 ,0.092, P = 0.42, N = 1101) | (0.5, 0.23, P = 0.21, N = 575) |
| rs767772939(CT)/7:128350744 | FAM71F1 | (0.096, 0.26, P = 4.1e-06, N = 1958) | (0.79, 0.01, P= 0.93, N = 651) | (0.82 ,-0.16, P = 0.29, N = 1101) | (0.8, 0.11, P = 0.66, N = 575) |
| rs17837468(A)/7:138309274 | SVOPL | (0.88, 0.0088, P = 3.8e-05, N = 7372) | (0.74, 0.03, P= 0.58, N = 651) | (0.76 ,-0.14, P = 0.3, N = 1101) | (0.74, 0.17, P = 0.43, N = 575) |
| rs17152416(T)/7:25765238 | AC003090.1 | (0.76, 0.0049, P = 3.9e-05, N = 7372) | (0.7, 0.03, P= 0.64, N = 651) | (0.71 ,-0.18, P = 0.17, N = 1101) | (0.69, 0.24, P = 0.25, N = 575) |
| rs4722699(T)/7:27456500* | intergenic | (0.27, -0.0048, P = 3.8e-05, N = 7372) | (0.23, -0.05, P= 0.49, N = 651) | (0.2 ,-0.0076, P = 0.96, N = 1101) | (0.22, 0.07, P = 0.73, N = 575) |
| rs1799211(T)/7:76240677 | UPK3B | (0.41, -0.0082, P = 4.2e-06, N = 1768) | (0.18, 0.04, P= 0.7, N = 401) | (0.24 ,0.037, P = 0.79, N = 1101) | (0.18, 0.31, P = 0.21, N = 325) |
| rs9314463(A)/8:2525166 | RP11-134O21.1 | (0.19, 0.27, P = 9.1e-07, N = 1958) | (0.45, -0.01, P= 0.83, N = 651) | (0.43 ,-0.073, P = 0.54, N = 1101) | (0.48, -0.05, P = 0.78, N = 575) |
| rs112508772(A)/9:140016354 | snoU13 | (0.025, 0.48, P = 7e-07, N = 1958) | (0.11, 0.14, P= 0.12, N = 651) | (0.056 ,-0.11, P = 0.67, N = 1101) | (0.13, 0.15, P = 0.58, N = 575) |
| rs7848854(C)/9:7766733* | intergenic | (0.11, 0.0083, P = 2.4e-05, N = 7372) | (0.02, -0.04, P= 0.85, N = 651) | (0.027 ,0.68, P = 0.06, N = 1101) | (0.02, -0.39, P = 0.58, N = 575) |
| $PTS_{51SNPs}$ | NA | NA | (NA, 0.021, P=0.60, N=651) | (NA, -0.056, P=0.12, N=1101) | (NA, -0.022, P=0.80, N=575) |
| $PTS_{27SNPs}$ | NA | NA | (NA, 0.025, P=0.53, N=651) | (NA, -0.081, P=0.021, N=1101) | (NA, 0.0036, P=0.97, N=575) |

**Supplementary Table 5. Mendelian randomization results for L-glutamine with sickle cell disease (SCD)-complications and estimated glomerular filtration rate (eGFR).** For complications, estimates are odds ratios (95% confidence intervals) for the effect of a 1 standard deviation increase in L-glutamine. For eGFR (0.07 mL/min per 1.172 $m^2$), estimates are effect size (standard error) for the effect of a 1 standard deviation increase in L-glutamine. All 51 genetic variants that are associated with L-glutamine at *P*<5x$10^{-5}$ are included in Model 1 analyses (29 SNPs from Shin et *al.*, 22 SNPs from Long et *al.*). In Model 2, we only kept 27 variants that were not pleiotropic (12 from Shin et *al.*, 15 from Long et *al.*). IVW: inverse variance-weighted. In light grey, we present MR replication results for L-glutamine and painful crises in the smaller GEN-MOD and OMG cohorts.

| Metabolite | Method | Painful crises (CSSCD) | Painful crises (GEN-MOD+OMG) | Cholecystectomy (CSSCD) | Retinopathy (CSSCD) | Leg ulcer (CSSCD) | Priapism (CSSCD) | Aseptic necrosis (CSSCD) | eGFR (CSSCD) |
|---|---|---|---|---|---|---|---|---|---|
| L-glutamine – Model 1 | IVW | 0.81 (0.63-1) P=0.086 | 0.77 (0.51- 1.16) P=0.21 | 0.94 (0.68-1.3) P=0.72 | 0.93 (0.67-1.3) P=0.66 | 1.1 (0.82-1.5) P=0.53 | 1 (0.78-1.4) P=0.81 | 0.88 (0.65-1.2) P=0.44 | 0.027 (0.057) P=0.64 |
| | MR-Egger | 0.76 (0.54-1.1) P=0.12 | 0.84 (0.5- 1.4) P=0.50 | 1.1 (0.76-1.7) P=0.52 | 0.85 (0.57-1.3) P=0.45 | 1.1 (0.75-1.6) P=0.66 | 1.1 (0.67-1.8) P=0.73 | 0.97 (0.65-1.4) P=0.88 | 0.061 (0.072) P=0.4 |
| | Weighted median | 0.77 (0.53-1.1) P=0.17 | 0.93 (0.55- 1.58) P=0.8 | 0.91 (0.58-1.4) P=0.67 | 0.82 (0.52-1.3) P=0.4 | 0.97 (0.62-1.5) P=0.88 | 1.1 (0.66-2) P=0.65 | 0.91 (0.59-1.4) P=0.67 | 0.015 (0.084) P=0.85 |
| L-glutamine – Model 2 | IVW | 0.68 (0.52-0.89) P=0.0048 | 0.82 (0.5- 1.34) P=0.44 | 0.84 (0.57-1.2) P=0.39 | 1.2 (0.78-1.7) P=0.46 | 1.2 (0.88-1.6) P=0.26 | 1.1 (0.78-1.5) P=0.6 | 0.85 (0.61-1.2) P=0.37 | -0.026 (0.079) P=0.74 |
| | MR-Egger | 0.74 (0.48-1.1) P=0.16 | 0.80 (0.42- 1.53) P=0.50 | 0.97 (0.59-1.6) P=0.92 | 1.1 (0.66-1.9) P=0.71 | 1.2 (0.75-1.9) P=0.46 | 1.2 (0.66-2.2) P=0.54 | 1 (0.64-1.6) P=0.96 | 0.012 (0.1) P=0.91 |
| | Weighted median | 0.73 (0.49-1.1) P=0.12 | 0.85 (0.44- 1.61) P=0.61 | 0.78 (0.46-1.3) P=0.36 | 1.2 (0.73-2.1) P=0.43 | 1.1 (0.66-1.8) P=0.72 | 1.2 (0.64-2.4) P=0.53 | 0.78 (0.46-1.3) P=0.37 | -0.0049 (0.099) P=0.96 |

**Supplementary Table 6. Pairwise association results between sickle cell disease (SCD) complications or estimated glomerular filtration rate (eGFR) and 129 profiled metabolites.** For associations with complications, we used logistic regression, with metabolites z-score-transformed and correction for age, sex, SCD genotypes, and hydroxyurea usage. Results for complications show odds ratio, 95% confidence interval (CI), and P-value. For eGFR, we performed linear regression with metabolites z-score corrected for SCD genotypes and hydroxyurea usage. Results for eGFR show beta-coefficient, standard error, and P-value. To account for the number of tests but also the correlation between metabolites, we derived empirical P-values using 100,000 permutations.

SEE EXCEL FILE

**Supplementary Table 7. Genetic associations between SNPs associated with 3-ureidopropionate and eGFR.** SNPs (effect alleles) are the lead variants in each gene region identified in a GWAS for 3-ureidopropionate levels in Europeans ($P$ <$5.0x10^{-5}$) (ref. [3]). Association effect size with eGFR in the CSSCD and GENMOD are beta-coefficient from linear regression in 0.07 mL/min per 1.172 $m^2$. *These variants were identified to be pleiotropic by Phenoscanner queries and were excluded from model 2 (**Methods**). At the bottom of the table, we also provide association results between polygenic trait scores (PTS) calculated using 22 3-ureidopropionate-associated SNPs ($PTS_{22SNPs}$) or after excluding pleiotropic variants ($PTS_{16SNPs}$). For the PTS, the effect size is per PTS standard deviation units.

| | | **Association result with 3-ureidopropionate -associated variants (effect allele frequency, effect size (beta), P-value, sample size** | | | |
|---|---|---|---|---|---|
| **SNP (hg19)** | **Gene region** | **3-ureidopropionate (Long et al)** | **3-ureidopropionate (Meta-analysis GEN-MOD+OMG)** | **eGFR (CSSCD)** | **eGFR (Meta-analysis GEN-MOD-OMG)** |
| rs75277555(T)/1:92235001 | TGFBR3 | (0.072, 0.16, P = 4e-06, N = 1956) | (0.057, -0.093, P = 0.43, N = 651) | (0.047 ,-0.008, P = 0.94, N = 859) | (0.06, -0.24, P = 0.15, N = 640) |
| rs8013355(C)/14:52871418 | intergenic | (0.32, 0.16, P = 1.7e-06, N = 1956) | (0.51, 0.012, P = 0.83, N = 651) | (0.48 ,-0.013, P = 0.78, N = 859) | (0.51, -0.05, P = 0.46, N = 640) |
| rs555045773(GA)/15:54319379 | UNC13C | (0.15, -1.1, P = 6.2e-06, N = 1956) | (0.23, -0.040, P = 0.58, N = 651) | (0.26 ,-0.069, P = 0.2, N = 859) | (0.23, 0.15, P = 0.09, N = 640) |
| rs59930743(C)/17:3393604 | ASPA | (0.22, 0.18, P = 2.8e-06, N = 1956) | (0.33, 0.030, P = 0.62, N = 651) | (0.29 ,0.013, P = 0.81, N = 859) | (0.33, 0.01, P = 0.87, N = 640) |
| rs56104151(T)/18:61129481 | intergenic | (0.17, 0.15, P = 8.1e-06, N = 1956) | (0.35, 0.064, P = 0.27, N = 651) | (0.35 ,0.031, P = 0.53, N = 859) | (0.34, 0.08, P = 0.32, N = 640) |
| rs78734409(A)/2:12335302 | AC096559.1 | (0.038, 0.51, P = 3e-06, N = 1956) | (0.026, 0.080, P = 0.63, N = 651) | (0.025 ,0.14, P = 0.37, N = 859) | (0.03, -0.36, P = 0.1, N = 640) |
| rs71394795(GTTTA)/2:12355843 | AC096559.1 | (0.034, 0.19, P = 4.9e-07, N = 1956) | (0.71, -0.024, P = 0.70, N = 651) | (0.66 ,0.016, P = 0.76, N = 859) | (0.71, 0.06, P = 0.49, N = 640) |
| rs13427576(T)/2:56523619 | CCDC85A | (0.12, 0.16, P = 9e-06, N = 1956) | (0.21, 0.036, P = 0.61, N = 651) | (0.22 ,0.0034, P = 0.95, N = 859) | (0.22, -0.1, P = 0.31, N = 640) |
| rs11704820(G)/22:24912248 | UPB1 | (0.43, 0.16, P = 4.3e-07, N = 1956) | (0.30, -0.14, P = 0.017, N = 651) | (0.32 ,0.0044, P = 0.93, N = 859) | (0.29, -0.01, P = 0.93, N = 640) |
| rs77020847(G)/3:150016629 | intergenic | (0.013, 0.24, P = 2.1e-06, N = 1956) | (0.015, -0.13, P = 0.60, N = 651) | (0.014 ,0.074, P = 0.71, N = 859) | (0.01, 0.65, P = 0.05, N = 640) |
| rs6788347(C)/3:37754694* | ITGA9 | (0.5, 0.19, P = 4e-06, N = 1956) | (0.53, 0.066, P = 0.25, N = 651) | (0.52 ,0.081, P = 0.1, N = 859) | (0.53, 0.06, P = 0.42, N = 640) |
| rs4698029(G)/4:10312798* | intergenic | (0.19, -0.39, P = 3.5e-17, N = 1956) | (0.23, -0.096, P = 0.17, N = 651) | (0.21 ,-0.033, P = 0.56, N = 859) | (0.22, 0.05, P = 0.56, N = 640) |
| rs13135526(A)/4:128932595* | C4orf29 | (0.51, -0.08, P = 9.1e-06, N = 1956) | (0.60, -0.088, P = 0.12, N = 651) | (0.61 ,0.057, P = 0.25, N = 859) | (0.6, 0.01, P = 0.92, N = 640) |
| rs2725772(T)/4:140438033* | SETD7 | (0.84, 0.21, P = 8.9e-08, N = 1956) | (0.50, -0.016, P = 0.78, N = 651) | (0.54 ,0.08, P = 0.093, N = 859) | (0.51, -0.05, P = 0.52, N = 640) |
| rs11735831(A)/4:9951591* | SLC2A9 | (0.22, -0.43, P = 1.4e-23, N = 1956) | (0.46, 0.054, P = 0.343, N = 651) | (0.45 ,-0.029, P = 0.54, N = 859) | (0.46, 0.06, P = 0.45, N = 640) |
| rs33379(C)/5:171099634* | intergenic | (0.44, 0.16, P = 4e-06, N = 1956) | (0.83, -0.011, P = 0.88, N = 651) | (0.77 ,-0.033, P = 0.58, N = 859) | (0.83, -0.1, P = 0.33, N = 640) |
| rs76129636(G)/6:130978833 | intergenic | (0.12, -0.41, P = 9.6e-06, N = 1956) | (0.24, -0.014, P = 0.83, N = 651) | (0.26 ,-0.0077, P = 0.89, N = 859) | (0.24, -0.13, P = 0.16, N = 640) |

| | | | | | |
|---|---|---|---|---|---|
| rs113133874(T)/6:33379903 | PHF1 | (0.11, -0.42, P = 7e-06, N = 1956) | (0.017, -0.062, P = 0.78, N = 651) | (0.021 ,-0.051, P = 0.76, N = 859) | (0.02, 0.15, P = 0.61, N = 640) |
| rs56286439(C)/7:14307105 | DGKB | (0.2, -0.13, P = 4.8e-06, N = 1956) | (0.83, -0.005, P = 0.95, N = 651) | (0.82 ,-0.0045, P = 0.94, N = 859) | (0.83, -0.03, P = 0.79, N = 640) |
| rs2352451(G)/8:112781984 | intergenic | (0.74, 0.18, P = 5.3e-06, N = 1956) | (0.75, 0.034, P = 0.59, N = 651) | (0.78 ,-0.073, P = 0.21, N = 859) | (0.75, -0.04, P = 0.67, N = 640) |
| rs7006208(A)/8:124157948 | TBC1D31 | (0.17, -0.96, P = 8e-06, N = 1956) | (0.25, 0.033, P = 0.61, N = 651) | (0.26 ,-0.087, P = 0.11, N = 859) | (0.25, 0.05, P = 0.56, N = 640) |
| rs74795659(A)/8:73633099 | KCNB2 | (0.068, 0.15, P = 9e-06, N = 1956) | (0.031, 0.13, P = 0.43, N = 651) | (0.023 ,0.24, P = 0.12, N = 859) | (0.03, -0.24, P = 0.28, N = 640) |
| $PTS_{22SNPs}$ | NA | NA | (NA, 0.013, P=0.73, N=651) | (NA, 0.069, P=0.044, N=859) | (NA, -0.017, P=0.065, N=649) |
| $PTS_{16SNPs}$ | NA | NA | (NA, 0.012, P=0.74, N=651) | (NA, 0.082, P=0.016, N=859) | (NA, -0.022, P=0.55, N=649) |

**Supplementary Table 8. Mendelian randomization results for 3-ureidopropionate with estimated glomerular filtration rate (eGFR) in sickle cell disease (SCD) patients.** Estimates are effect size (standard error) in eGFR units (0.07 mL/min per 1.172 $m^2$) for the effect of a one standard deviation increase in genetically-controlled 3-ureidopropionate. All 22 genetic variants that are associated with 3-ureidopropionate at $P<5x10^{-5}$ are included in Model 1. In Model 2, we only kept 16 variants that were not pleiotropic. IVW: inverse variance-weighted. In light grey, we present MR replication results for 3-ureidopropionate and eGFR in the smaller GEN-MOD and OMG cohorts.

| Metabolite | Method | eGFR (CSSCD) | eGFR (GEN-MOD) | eGFR (OMG) |
|---|---|---|---|---|
| 3-ureidopropionate – Model 1 | IVW | 0.078 (0.023) P=0.00087 | -0.093 (0.059) P=0.12 | -0.076 (0.059) P=0.19 |
| | MR-Egger | 0.089 (0.048) P=0.077 | -0.16 (0.089) P=0.082 | -0.024 (0.090) P=0.79 |
| | Weighted median | 0.068 (0.045) P=0.13 | -0.13 (0.076) P=0.65 | 0.027 (0.096) P=0.77 |
| 3-ureidopropionate – Model 2 | IVW | 0.07 (0.021) P=0.00097 | -0.082 (0.068) P=0.22 | -0.074 (0.068) P=0.28 |
| | MR-Egger | 0.088 (0.05) P=0.1 | -0.14 (0.099) P=0.17 | -0.057 (0.01) P=0.58 |
| | Weighted median | 0.068 (0.049) P=0.16 | -0.12 (0.081) P=0.15 | 0.077 (0.11) P=0.49 |

**Supplementary Figure 1.** Principal component analysis (PCA) of the metabolomic data before (**A**) and after (**B**) batch-effect correction using comBAT. Although the 3 different batches are clearly distinguishable before correction, comBAT pre-processing removes this effect. In each plot, the x- and y-axis represent the first and second principal components. The legend is the same for both plots.

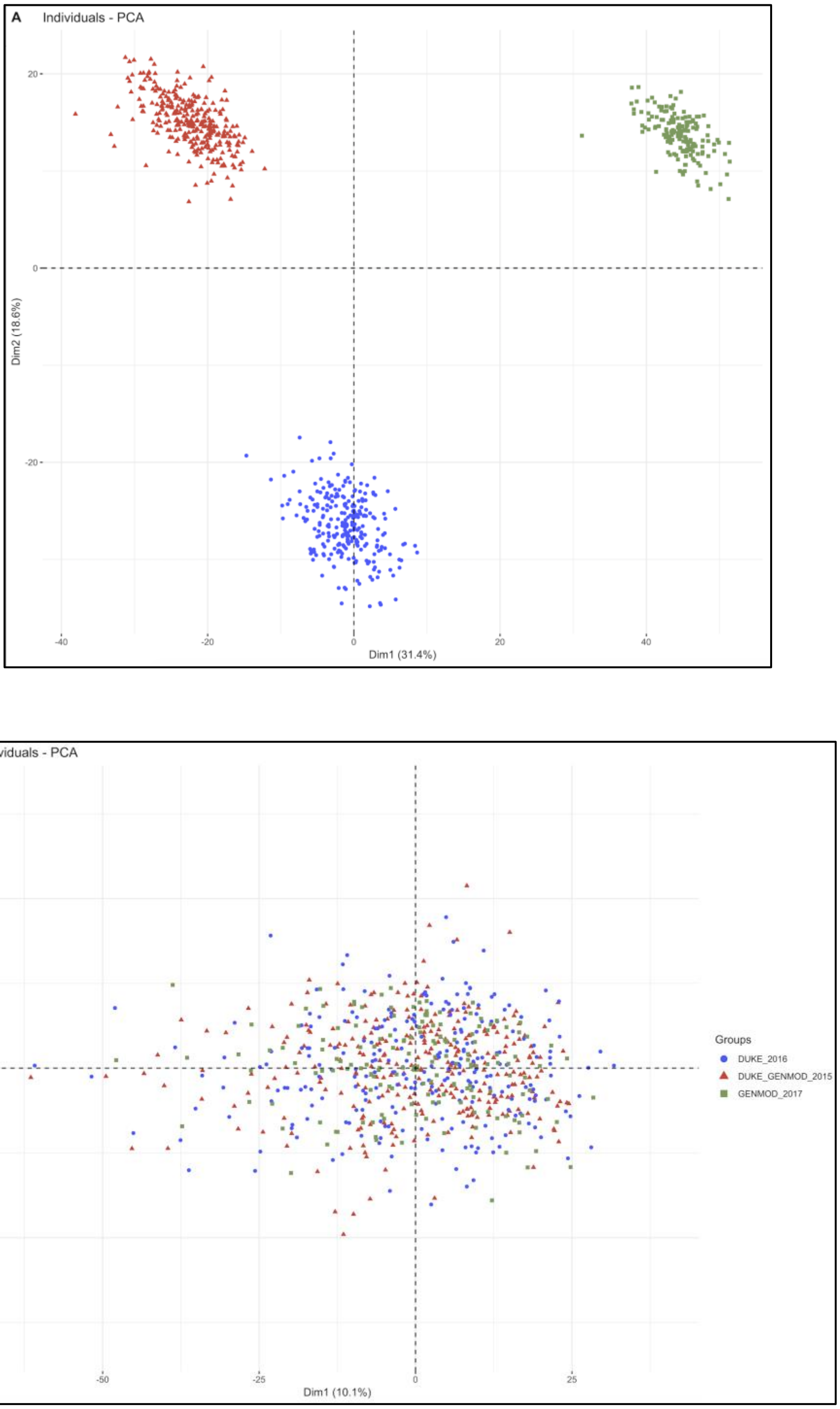

**Supplementary Figure 2.** 129 known metabolites grouped in super-pathways (**A**) and pathways (**B**) based on criteria from the human metabolome database (HMDB).

**A**

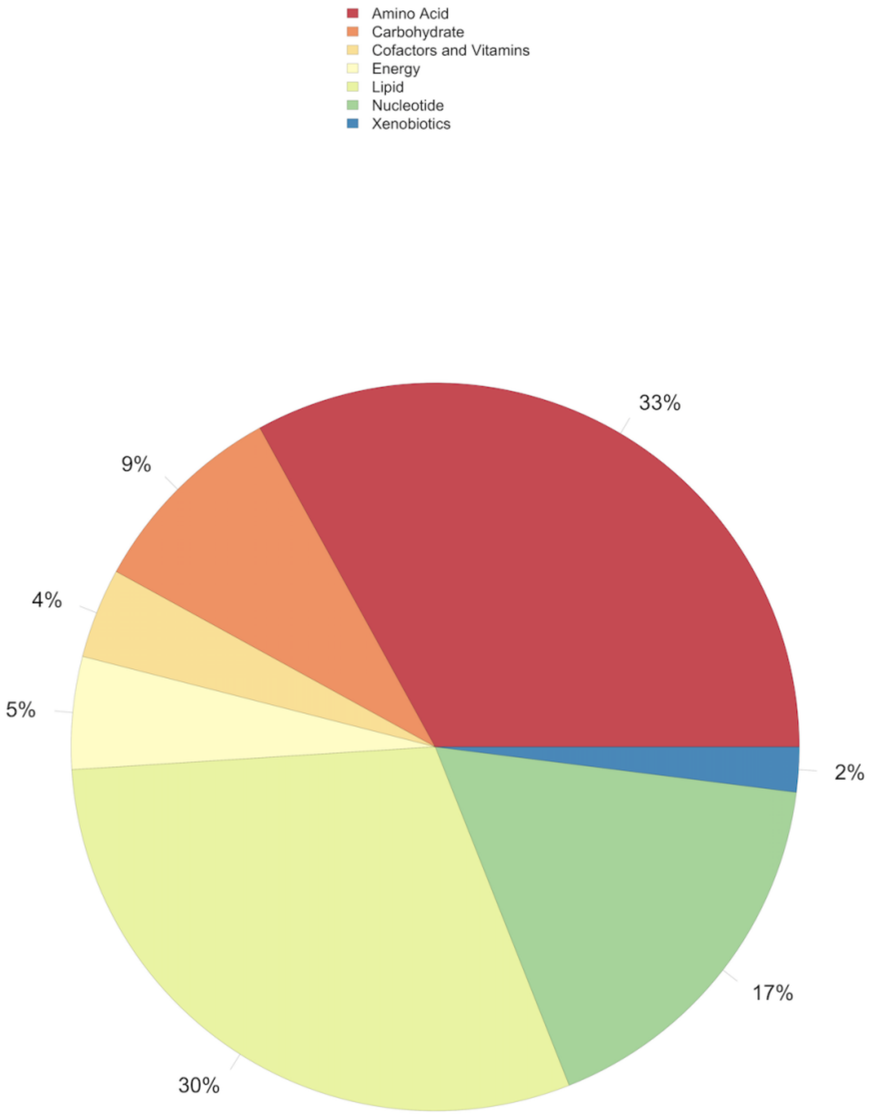

B

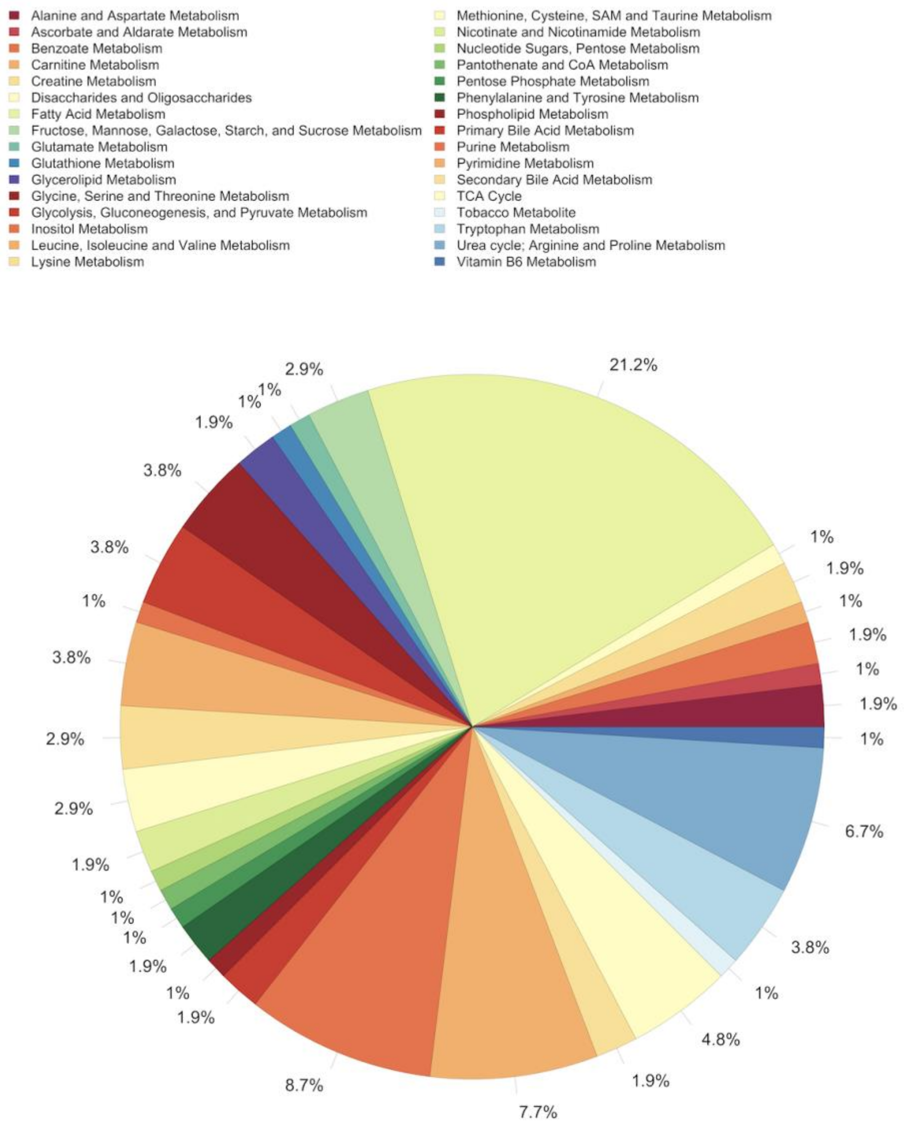
Alanine and Aspartate Metabolism
Ascorbate and Aldarate Metabolism
Benzoate Metabolism
Carnitine Metabolism
Creatine Metabolism
Disaccharides and Oligosaccharides
Fatty Acid Metabolism
Fructose, Mannose, Galactose, Starch, and Sucrose Metabolism
Glutamate Metabolism
Glutathione Metabolism
Glycerolipid Metabolism
Glycine, Serine and Threonine Metabolism
Glycolysis, Gluconeogenesis, and Pyruvate Metabolism
Inositol Metabolism
Leucine, Isoleucine and Valine Metabolism
Lysine Metabolism
Methionine, Cysteine, SAM and Taurine Metabolism
Nicotinate and Nicotinamide Metabolism
Nucleotide Sugars, Pentose Metabolism
Pantothenate and CoA Metabolism
Pentose Phosphate Metabolism
Phenylalanine and Tyrosine Metabolism
Phospholipid Metabolism
Primary Bile Acid Metabolism
Purine Metabolism
Pyrimidine Metabolism
Secondary Bile Acid Metabolism
TCA Cycle
Tobacco Metabolite
Tryptophan Metabolism
Urea cycle; Arginine and Proline Metabolism
Vitamin B6 Metabolism
21.2%
1%
1.9%
1%
1.9%
1%
1.9%
1%
6.7%
3.8%
1%
4.8%
1.9%
7.7%
8.7%
1.9%
1%
1.9%
1%
1%
1%
1.9%
2.9%
2.9%
3.8%
1%
3.8%
3.8%
1.9%
1%
1%
2.9%
